# The probability of edge existence due to node degree: a baseline for network-based predictions

**DOI:** 10.1101/2023.01.05.522939

**Authors:** Michael Zietz, Daniel S. Himmelstein, Kyle Kloster, Christopher Williams, Michael W. Nagle, Casey S. Greene

## Abstract

Important tasks in biomedical discovery such as predicting gene functions, gene-disease associations, and drug repurposing opportunities are often framed as network edge prediction. The number of edges connecting to a node, termed degree, can vary greatly across nodes in real biomedical networks, and the distribution of degrees varies between networks. If degree strongly influences edge prediction, then imbalance or bias in the distribution of degrees could lead to nonspecific or misleading predictions. We introduce a network permutation framework to quantify the effects of node degree on edge prediction. Our framework decomposes performance into the proportions attributable to degree and the network’s specific connections. We discover that performance attributable to factors other than degree is often only a small portion of overall performance. Degree’s predictive performance diminishes when the networks used for training and testing—despite measuring the same biological relationships—were generated using distinct techniques and hence have large differences in degree distribution. We introduce the permutation-derived edge prior as the probability that an edge exists based only on degree. The edge prior shows excellent discrimination and calibration for 20 biomedical networks (16 bipartite, 3 undirected, 1 directed), with AUROCs frequently exceeding 0.85. Researchers seeking to predict new or missing edges in biological networks should use the edge prior as a baseline to identify the fraction of performance that is nonspecific because of degree. We released our methods as an open-source Python package (https://github.com/hetio/xswap/).

## Introduction

Networks contain information about relationships between entities (referred to here as “edges” between “nodes”). A node’s degree is the number of edges it has in the network. Networks contain many nodes, whose degrees can be aggregated to form the network’s degree distribution. Because different nodes can have very different degrees, real networks have a variety of degree distributions (Figure 1), and they commonly exhibit degree imbalance [1,2,3,4]. This is especially true for networks encoding biomedical knowledge or assays, where natural forces such as preferential attachment inherent to the problem domain combine with observation-based influences such as study methodology to create non-uniform degree distributions (Figure 1).

**Figure 1:**
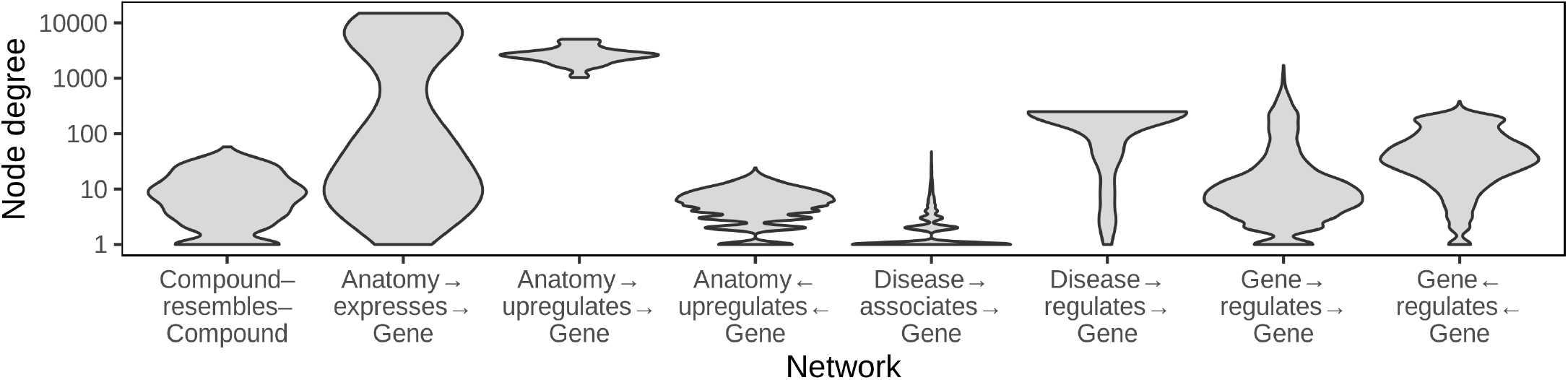
Biomedical networks are characterized by non-uniform degree distributions. Eight degree distributions are plotted for six edge types Hetionet v1.0 [5]. Hetionet integrates subnetworks for 24 different edge types, the degree distributions of which are analyzed separately. Furthermore, bipartite (e.g. Anatomy→ expresses→ Gene) and directed (e.g. Gene→ regulates→ Gene) graphs (Hetionet edge types) have both source and target degrees that must be assessed separately. Undirected edge types (e.g Compound–resembles–Compound) have only a single degree distribution. Degree distributions are non-uniform and vary greatly between different networks. The y-axis is log_10_-scaled to accommodate the common occurrence where most nodes have low degree while a small portion of nodes have high degree. Several distributions have nodes that reach the maximum degree, corresponding to a node being connected to all other possible nodes. Zero-degree nodes are not displayed, since methodological limitations often result in edge data only existing for a subset of nodes.

Degree is an important metric for differentiating between nodes, and it appears in many common edge prediction features [6]. However, reliance on degree can pose problems for edge prediction. First, bias in networks can distort node degree so that a difference in degree between two nodes in a given network may not reflect a true difference in number of relationships. Second, edge prediction methods that rely heavily on degree may be nonspecific—predicting trivial rather than insightful new relationships.

Most biomedical networks are imperfect representations of the true set of relationships. Real networks often mistakenly include edges that do not exist and exclude edges that do exist. How well a network represents the true relationships it attempts to represent depends on a number of factors, especially the methods used to generate the data in the network [7,8,9]. We define “degree bias” as the type of misrepresentation that occurs when the fraction of incorrectly existent/nonexistent relationships depends on a node’s degree. Depending on the type of data being represented, degree biases can arise due to experimental methods, inspection bias, or other factors [7].

Inspection bias indicates that entities are not uniformly studied [10], and it is likely to cause degree bias when networks are constructed using hypothesis-driven findings extracted from the literature, as newly-discovered relationships are not randomly sampled from the set of all true relationships. Though there is a high correlation between the number of publications mentioning a gene and its degree in low-throughput interaction networks, the number of publications mentioning a gene has little correlation with its degree in a systematically-derived protein interaction network [11]. This suggests that many poorly connected genes in non-systematic protein interaction networks are due to inspection bias, i.e. a lack of study, rather than a lack of biological function. For networks with a large inspection bias, reliance on degree can lead to predictions that have good metrics when assessed by cross validation but little ability to generalize.

Another reason why a reliance on degree can be unfavorable is that degree imbalance can lead to prediction nonspecificity. Nonspecific predictions are not made on the basis of the specific connectivity information contained in a network. For example, Gillis et al. examined the concept of prediction specificity in the context of gene function prediction and found that many predictions appear to rely primarily on multifunctionality and could be “potentially misleading with respect to causality” [12]. Degree imbalance leads high-degree nodes to dominate in the predictions made by degree-associated methods [13], which are effective predictors of connections in some biological networks [14]. Consequently, degree-based predictions are more likely nonspecific, meaning the same set of predictions performs well for different tasks.

Depending on the prediction task, edge predictions involving very high degree nodes may be undesired, uninsightful, or nonspecific. While predictions based primarily on degree may be acceptable for some tasks, generating less obvious insights from networks requires drawing inferences from the specific connections and network structure between nodes. Model evaluation is challenging in this context: nonspecific or trivial predictions can dominate performance evaluations and may actually be correct, even if they are not the desired outputs of the predictive model. For example, predicting that the highest degree node in a network shares edges with the remaining nodes to which it is not connected will often lead to many correct predictions, despite this prediction being generic to all other nodes in the network.

Degree is important in edge prediction, but it can cause undesired effects. Degree-based features should often be included in the interpretation of predictions to disentangle desired from non-desired effects and to effectively evaluate and compare predictive models. We sought to directly measure the effect of node degree on edge prediction methods. We introduce a permutation-based framework and software implementation to find edge existence probabilities due to node degree and to quantify the contribution of degree to edge prediction methods. This method allows edge predictions to be evaluated in the context of degree and its effects on the prediction task. Our results demonstrate that degree-associated methods are very effective for reconstructing a network using a subsampled holdout. However, these methods are ineffective for predicting edges between networks measuring the same biological processes in targeted and systematic ways because such networks have distinct degree distributions. Using multiple different networks, we provide evidence that degree has a strong effect on the probability of edge existence and that our permutation-based edge prior best quantifies this probability.

## Methods

### Network permutation

Network permutation is a way to produce new networks by randomizing the connections of an existing network. Specialized permutation strategies can be devised that randomize some aspects of networks while retaining other features. Comparing between permuted and unpermuted networks gives insight to the effects of the retained network features. For example, an edge prediction method that has superior reconstruction performance on a network compared to its permutations likely relies on information that is eliminated by permutation. Conversely, identical predictive performance on true and permuted networks indicates that a method relies on information that is preserved during permutation.

Network permutation is a flexible framework for analyzing other methods, because it generates complete networks that can be analyzed independently. We use network permutation to isolate degree and determine its effects in different contexts. Degree-preserving network permutation obscures true connections and higher-order connectivity information (e.g., community structure), while retaining node degree, and, thereby, the network’s degree sequence. Thanks to the flexibility of permutation, our framework can quantify the effect of degree on any network edge prediction method.

### XSwap algorithm

Hanhijärvi, et al. presented XSwap [15], an algorithm for the randomization (“permutation”) of unweighted networks (Figure 2A). The algorithm picks two existing edges at random ({ab, cd}) and—if the edges constitute a valid swap—exchanges the targets between the edges ({ad, cb}; Supplemental Table 1). This process is repeated a user-specified number of times. In general, the number of exchanges should be chosen to be sufficiently large that the fraction of original edges retained in the permuted network is near its asymptotic value as the number of exchanges increases to infinity. The asymptotic fraction of original edges retained in permutation depends on network density, and higher density networks require more swap attempts per edge to reach their asymptotic fraction (Figure 9).

**Figure 2:**
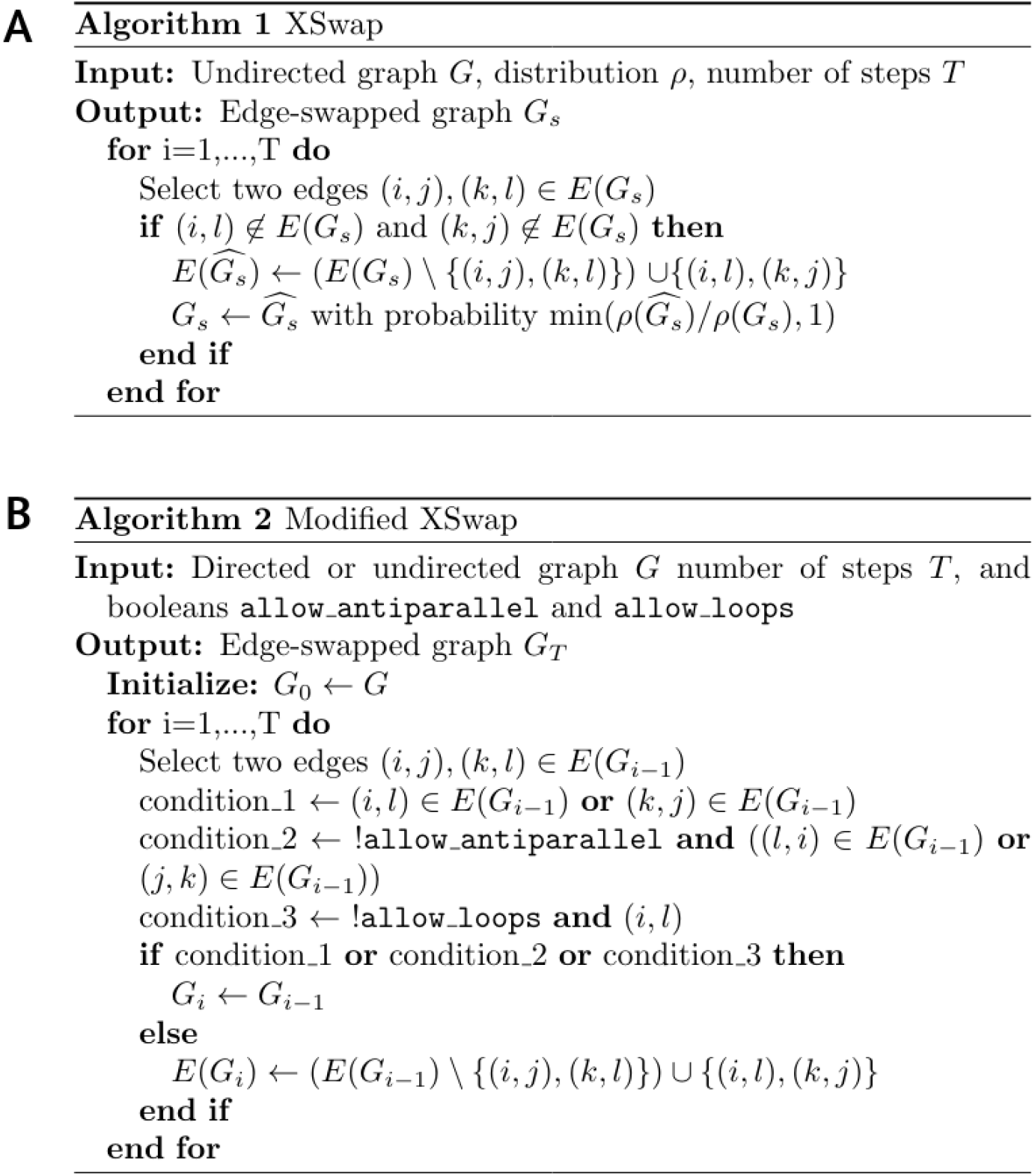
XSwap algorithm pseudocode. **A**. XSwap algorithm presented by Hanhijärvi, et al. [15]. **B**. Extension of the XSwap algorithm to other types of networks.

We modified the original XSwap algorithm by adding two parameters, `allow_loops` (a-a), and `allow_antiparallel` (a-b and b-a) that allow a greater variety of network types to be permuted (Figure 2B and Supplemental Table 1). The motivation for these generalizations is to make the permutation method applicable both to directed and undirected graphs, as well as to networks with different types of nodes, variously called multipartite, heterogeneous, or multimodal networks. Specifically, in the modified algorithm two chosen edges constitute a valid swap if they preserve degree for all four involved nodes and do not violate the user-specified parameters.

When permuting bipartite networks, our method ensures that each node’s class membership and within-class degree is preserved. Similarly, heterogeneous networks should be permuted by considering each edge type as a separate network [16,17]. This way, each node retains its within-edge-type degree for all edge types. We provide documentation for parameter choices depending on the type of network being permuted in the GitHub repository (https://github.com/hetio/xswap). The original algorithm and our proposed modification are given in Figure 2.

### Edge prior

We introduce the edge prior to quantify the probability that two nodes are connected based only on their degree. The edge prior can be estimated using the fraction of permuted networks in which a given edge exists—the maximum likelihood estimate for the binomial distribution success probability. Based only on permuted networks, the edge prior does not contain any information about the true edges in the (unpermuted) network. The edge prior is a numerical value that can be computed for every pair of nodes that could potentially share an edge; we compared its ability to predict edges in three tasks, discussed in prediction tasks.

### Analytical approximation of the edge prior

Because network permutation can be computationally intensive, we also considered whether the probability of an edge existing across permuted networks has a simple closed-form expression. We were unable to find a closed-form solution giving the edge prior without assuming that the probability of any given edge existing is independent of all other potential edges, which we believe is not valid for XSwap. Nonetheless, we discovered a good analytical approximation to the edge prior that is particularly good for networks with many nodes and fewer edges (Figure 3). Let *m* be the total number of edges in the network, and *u*_*i*_, *v*_*j*_ be the source and target degrees of a node pair, respectively. An approximation of the edge prior is

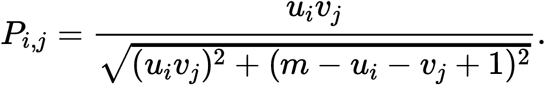

**Figure 3:**
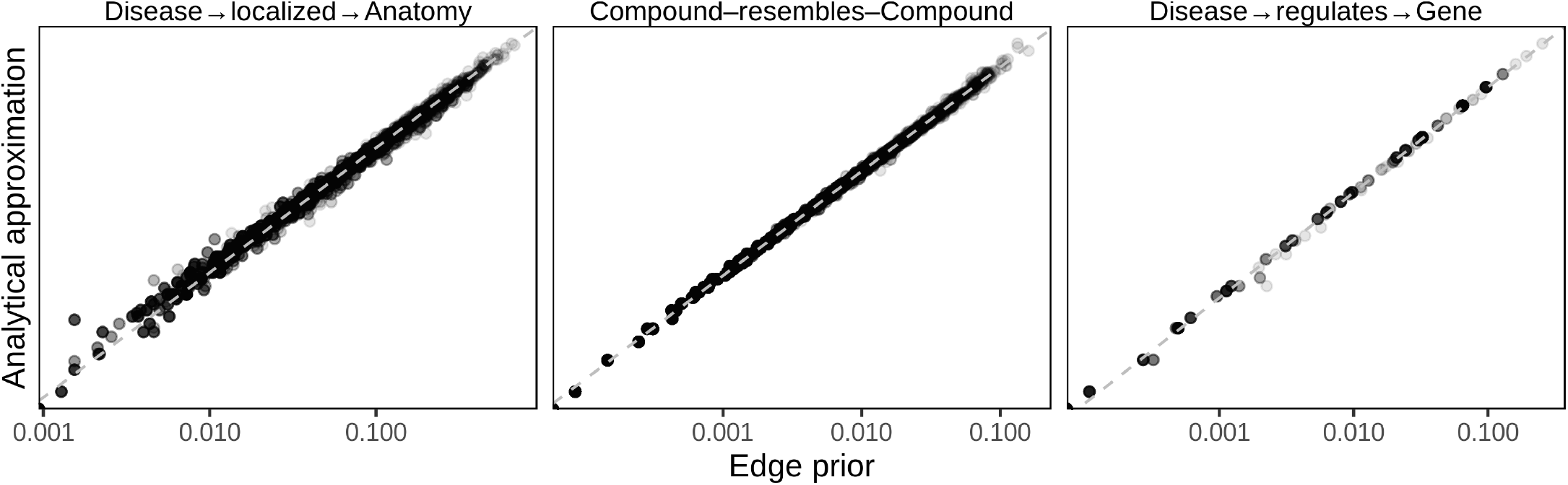
The XSwap-derived edge prior can be analytically approximated. The analytical approximation is plotted against the XSwap-derived edge prior for three networks (edge types) from Hetionet. The strong correlation suggests that the approximation will be suitable for applications where computation time is a limiting factor.

Further discussion of this approximate edge prior and an derivation are available in the supplement.

### Prediction tasks

We performed three prediction tasks to assess the performance of the edge prior. We compared the permutation-based prior with two additional predictors: our analytical approximation of the edge prior and the product of source and target degree, scaled to the range [0, 1] so that we could assess its calibration as well as its discrimination. We used 20 biomedical networks from the Hetionet heterogeneous network [5] that had at least 2000 edges for the first two tasks (Supplemental table). In the first task, we computed the degree-based predictors (edge prior, scaled degree product, and analytical prior approximation), and predicted the original edges in the network by rank-ordering node pair edge predictions by the node pairs’ predictor values. We used node pairs that lacked an edge in the original network as negative examples and those with an edge as positive examples. To assess the methods’ predictive performances, we computed the area under the receiver operating characteristic (AUROC) curve for all three predictors. In the second task, we sampled 70% of edges from each of the networks, computed predictors on the sampled network, then predicted held-out edges. For this task, negative examples were node pairs in which an edge did not exist in either original or sampled network, while positive samples were those node pairs without an edge in the sampled network but with an edge in the original network.

The third task evaluated the ability of the edge prior to generalize to new degree distributions. We used two domains where networks were available which shared nodes but had different degree distributions. Protein-protein interactions (PPI) and transcription factor-target gene (TF-TG) relationships had networks created both by literature curation of low-throughput, hypothesis-driven research and by high-throughput, systematic, hypothesis-free experimentation. For the PPI networks, we used the STRING network, which incorporates literature-mining to find relationships [18] and a combination of the high-throughput, proteome-scale interaction networks from Rual et al. [10] and Rolland et al. [11]. We used a transcription factor-target gene (TF-TG) literature-derived network from Han et al. [19] and a high-throughput network from Lachmann et al. [20]. The pairs of networks for PPI and TF-TG data sources are ideal because in one we expect inspection bias and in the other we do not.

As a further basis of comparison, we added a time-resolved co-authorship network, which we partitioned by time to create two separate networks. We created the co-authorship network of bioRxiv bioinformatics preprints using the Rxivist [21,22] database, which was generated by crawling the bioRxiv server. Unlike the other two networks, co-authorship does not have degree bias, as the network faithfully represents all true co-author relationships. We include this network to offer a comparative prediction task in which the degree distributions between training (posted before 2018) and testing (posted during or after 2018) are not dramatically different (Figure 4A). The goal of the third prediction task is to determine predictor generalizability for network reconstruction between different degree distributions, especially predicting a network without degree bias using predictors from a degree-biased network. Further information about the networks used can be found in the supplement.

**Figure 4:**
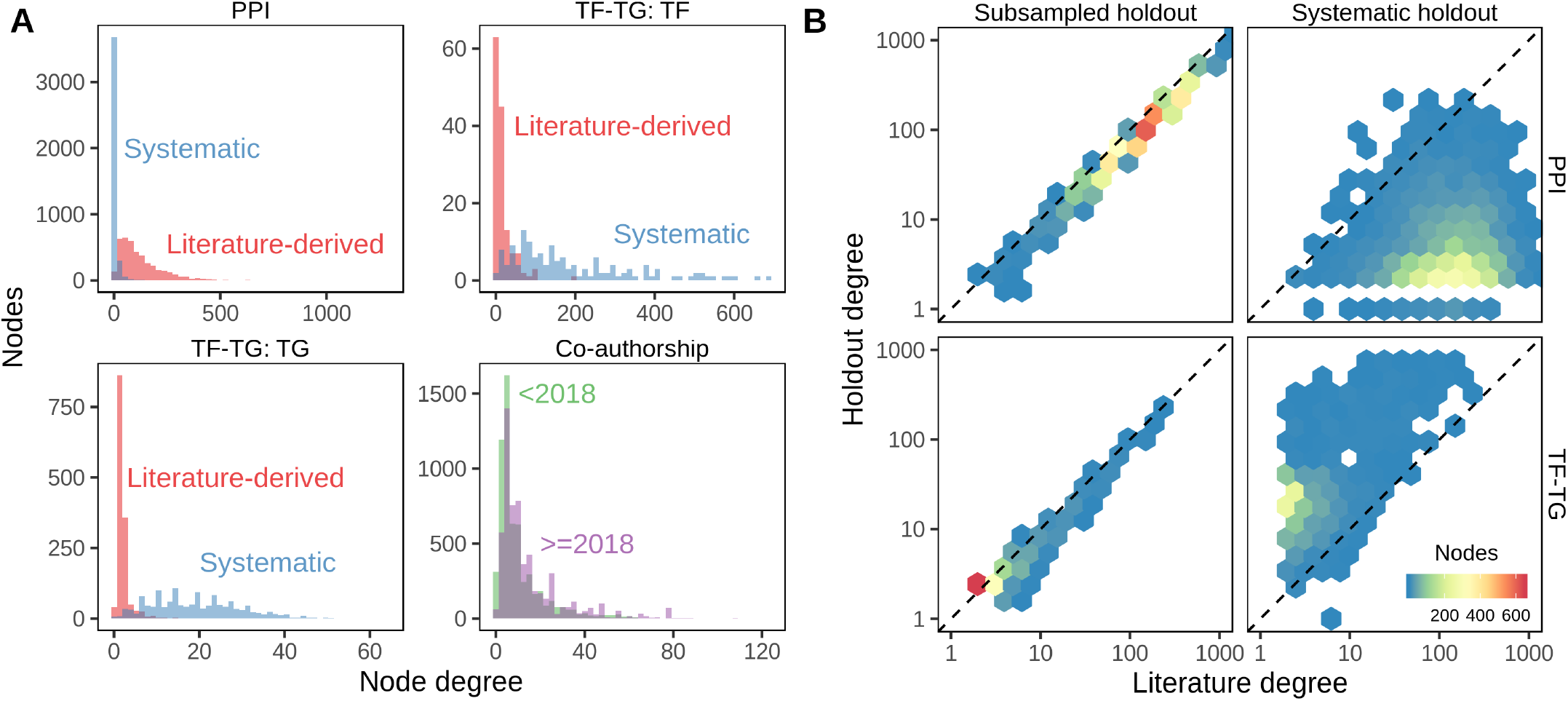
**A**. Degree distributions of networks with and without degree bias can be very different. Data on PPI and TF-TG were split between literature-derived and systematically-derived networks. In both cases, the networks exhibit large differences in degree distribution. Co-authorship relationship networks split by date of first co-authorship roughly share their degree distributions. **B**. Comparison of individual node degrees between different networks. Not only are the overall degree distributions different, but individual nodes can have systematically different degrees between two networks. Uniform random sampling produces linearly-correlated node degree, while non-random sampling produces non-correlated degree. Systematically-derived networks are not uniformly sampled from literature-derived networks or vice versa. 70% of literature edges were sampled with uniform probability for the “Subsampled holdout” network.

### Degree-grouping

Our method for degree-preserving permutation produces randomized networks that share few of their edges with the original network. The predictor values for two node pairs with the same source and target degree are drawn from the same distribution in permuted networks, so nodes with equal degree can be grouped when summarizing predictors. For a given node pair, degree grouping treats other node pairs with the same degrees as additional permutations [23]. We used this strategy to augment the number of predictor values for each node pair in permuted networks, allowing node pairs to have more permuted predictor values than permuted networks. Degree grouping greatly increased the effective number of permutations for nodes with frequently observed degrees. We used degree grouping throughout our analyses.

### Implementation and source code

We implemented our modified version of the XSwap algorithm as an open-source Python package. The package contains modules for permuting networks, computing the edge prior, and converting networks between adjacency matrix and edge list formats. Additionally, we include the analytical approximation of the edge prior and functionality to assign unique identifiers to nodes. The Python package is available on the Python Packaging Index under the name “xswap”. The full source code is freely available under the BSD 2-Clause License (https://github.com/hetio/xswap).

The edge swap mechanism—implemented in C++ for greater speed—uses a bitset to avoid producing edges which violate the conditions for a valid swap. While the full bitset implementation is faster for smaller networks, our package uses a compressed bitset [24] when a network would occupy memory above a user-adjustable threshold. In addition to the validity conditions already described, our package allows specific edges to be excluded from permutation, and every network permutation returns both a permuted network and summary information about the numbers of swaps attempted, performed, and the reasons why invalid swaps were rejected.

In addition to the Python package, all code to generate the analyses and figures is available at https://github.com/greenelab/xswap-analysis. The manuscript was written using the Manubot software [25], which allows anyone to provide feedback or modifications via the public repository at https://github.com/greenelab/xswap-manuscript.

## Results

### Node degree bias is prevalent

We found examples of node degree bias in the PPI and TF-TG networks we investigated. Figure 4 shows node degree in separate networks for the same type of data. For the PPI networks, the literature-derived network has a larger mean degree and a longer tail than the systematic network, while in the TF-TG networks this relationship is reversed. Because the TF-TG network contained far more transcription factors than target genes (144 and 1406, respectively), the distributions of target degrees were far more compact than those of source degrees. Unlike the PPI and TF-TG networks, the co-authorship networks were split by date of first co-authorship and did not exhibit a great difference in their degree distributions. All three types of networks (PPI, TF-TG, and co-authorship) exhibit degree imbalance to varying extents. These results indicate that, depending on the methods by which the represented data were generated, networks of the same type of data may have overall degree distributions that differ greatly (Figure 4A), and they may even assign very different degree to the same nodes (Figure 4B).

### The edge prior encapsulates degree

In the first prediction task, we computed three predictors—the XSwap edge prior, an analytical approximation to the edge prior, and the (scaled) product of source and target node degree—on networks from Hetionet. We then evaluated the extent to which these predictors—treated as predictions themselves—could reconstruct the 20 networks (Supplemental table). The XSwap-derived edge prior reconstructed many of the networks with a high level of performance, as measured by the AUROC. Of the 20 individual networks we extracted from Hetionet, 17 had an edge prior self-reconstruction AUROC >= 0.95, with the highest reconstruction AUROC at 0.9971 (network was the Compound–downregulates–Gene edge type). Meanwhile, the lowest self-reconstruction performance (AUROC = 0.7697) occurred in the network having the fewest node pairs (network was the Disease– localizes–Anatomy edge type).

The three predictors that we compared were highly correlated (Spearman rank correlation over 0.984 for all 20 networks). The three predictors also had very similar AUROC reconstruction performance values for the first, second, and third prediction tasks (max difference < 0.027) because AUROC is rank-based. The edge prior was slightly better than the approximations in 12 of 20 networks. However, while the AUROC results were similar, the predictors were very different in their levels of calibration—the ability of the model to correctly estimate edge existence probabilities. The edge prior was very well calibrated for all networks in the first and second tasks, and it provided the best calibration of the three predictors for each of the prediction tasks (Figure 6A). As the edge prior was not based on the networks’ true edges, these results indicated that degree sequence alone was highly informative and that permutation was the only approach in our comparison that provided a well-calibrated model.

**Figure 5:**
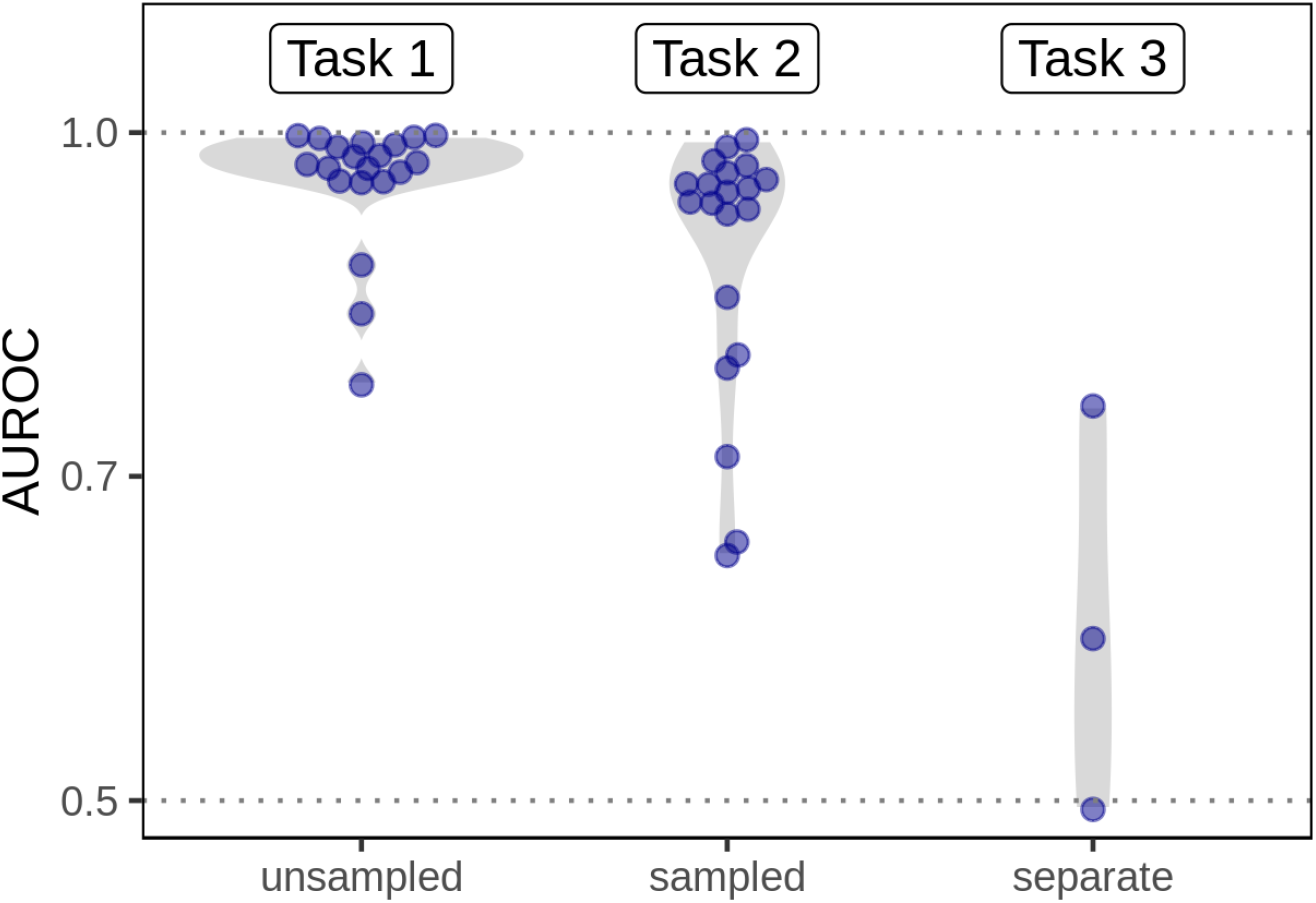
Degree can predict edges within a given network but does not generalize to networks with diferent degree distributions. The edge prior is able to reconstruct the networks on which it was computed (Task 1, “unsampled”, 20 different networks) with high performance. When computed on a sampled network, the edge prior can reconstruct the unsampled network with slightly lower performance (Task 2, “sampled”, 20 different networks). However, when computed on a completely different network (having a different degree distribution) of the same type of data, the edge prior’s performance is greatly reduced (Task 3, “separate”, 3 different networks). The performance reduction from computing predictors on sampled networks is real but far smaller compared to a new degree distribution. This indicates that while degree can be effective for network reconstruction, it is far less effective in predicting edges from a different degree distribution.

**Figure 6:**
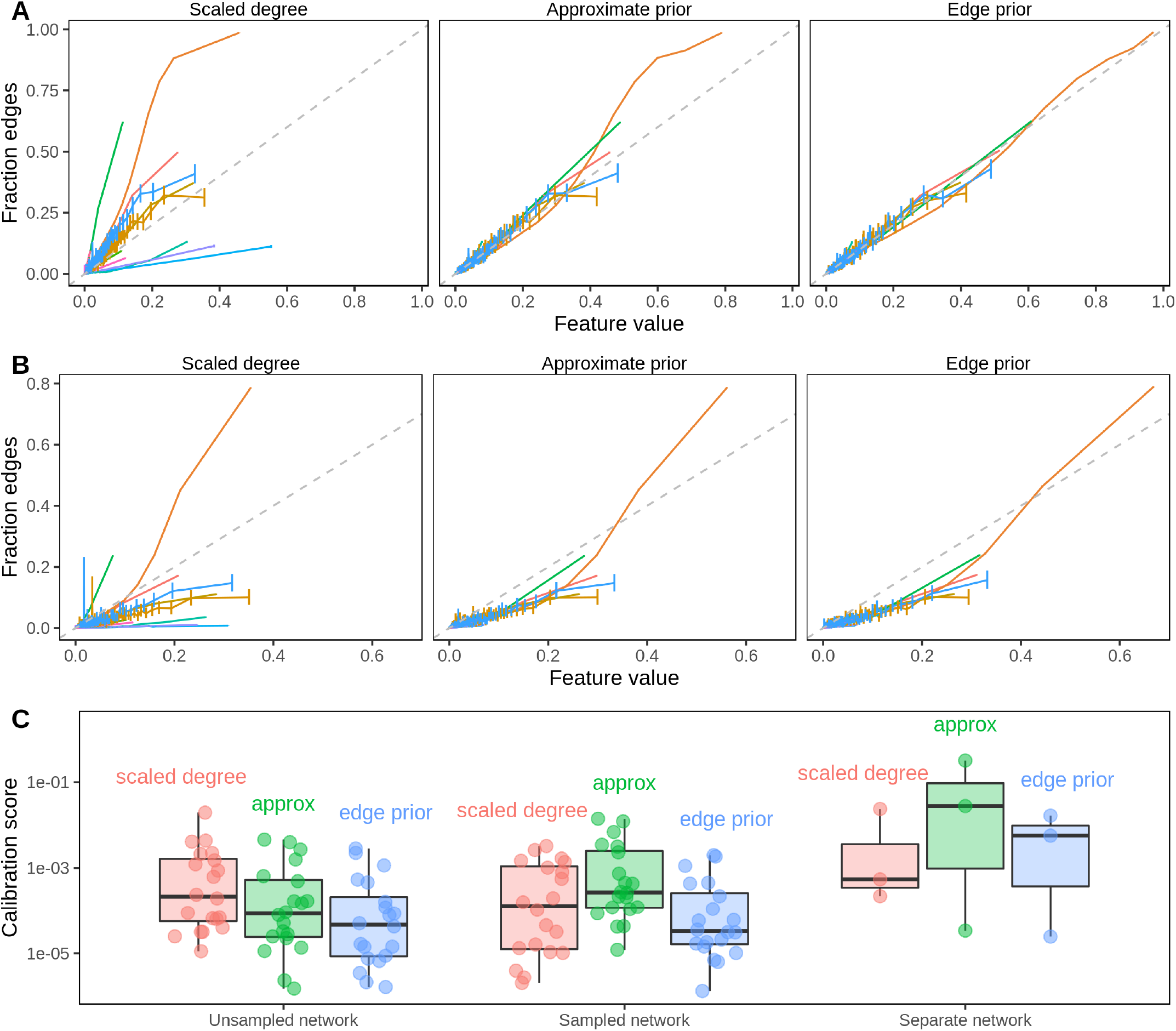
The edge prior accurately assigns the probability of edge existence. **A**. Calibration curves for full network reconstruction of 20 networks from Hetionet. For every unique predictor value on the horizontal axis, the fraction of node pairs with that predictor value having an edge in the network is shown on the vertical axis. The permutation-based edge prior’s calibration was superior to the other two strategies based on degree. **B**. Calibration curves for sampled network reconstruction. The edge prior shows superior calibration in the 20 Hetionet networks. **C**. Individual Hetionet edge type calibration estimated by the two-component decomposition of the Brier score, in which lower scores indicate better calibration. The edge prior has excellent calibration in unsampled and sampled networks, and each considered method is sensitive to shifts in the degree distribution.

The second prediction task mirrored the first task, but it involved reconstructing networks based on subsampled networks with only 70% of the original edges. Because edges were sampled uniformly without replacement, the subsampled networks share similar degree distributions to the original networks (see Figure 4B). Unlike in the first task, edges that were present in the sampled network were not tested and therefore are not included in the performance metrics. The results of the second prediction task further demonstrate a high level of performance for degree-sequence-based node pair predictors (Figure 5). The edge prior was able to reconstruct the unsampled network with an AUROC of greater than 0.9 in 14 of 20 networks. As was observed in the first task, node pair predictors computed in the second task were highly rank-correlated, meaning the AUROC values for different predictors were similar. While performance was slightly lower in the second task than the first, many networks were still well-reconstructed. The edge prior was the best calibrated predictor for both tasks.

In the third prediction task, we computed the three edge predictors for paired networks representing data from PPI, TF-TG, and bioRxiv bioinformatics pre-print co-authorship. The goal of the task was to compare predictive performance across different degree distributions for the same type of data. We find that the task of predicting systematically-derived edges using a network with degree bias is significantly more challenging than network reconstruction, and we find consistently lower performance compared to the other tasks (Figure 5). The edge prior was not able to predict the separate PPI network better than by random guessing (AUROC of roughly 0.5). Only slightly better was its performance in predicting the separate TF-TG network, at an AUROC of 0.59. We find superior performance in predicting the co-authorship relationships (AUROC 0.75), which was expected as the network being predicted shared roughly the same degree distribution as the network on which the edge prior was computed. The results of the third prediction task show that a difference in degree distribution between the network on which predictors are computed and the network to be predicted can make prediction significantly more challenging.

The edge prior can be considered a baseline edge predictor that accurately captures degree’s contribution to the probability of an edge existing. The edge prior’s low performance in the third task indicates that degree is less helpful for edge prediction tasks in which training and testing networks do not share their degree distributions. Many biomedical prediction tasks can be framed as edge prediction tasks between different degree distributions. In drug repurposing, for example, existing compound-disease treatment relationships are unlikely to be randomly sampled from all true treatment relationships. However, all treatment relationships between existing compounds and diseases are desirable outputs in prediction. Edge predictions can be based on both underlying biological properties and network degree distributions. However, predictions based on biological properties may be more consistent and generalizable than those based on degree. Degree’s influence on edge prediction accuracy measures can reveal the relative contributions of these two factors.

### Degree can underly a large fraction of performance

We conducted a further edge prediction task as an example application of the edge prior and our permutation framework. To begin, we chose the STRING PPI network for the comparison and computed five edge prediction features (Supplemental table 2). The goal of the task was to reconstruct the network on which the features were computed. All five features were correlated with degree (Figure 7), which we quantified for a node pair using the product of source and target degrees. We expected features based on degree to show strong performance for a network reconstruction task without holdout, as found in the first prediction task.

**Figure 7:**
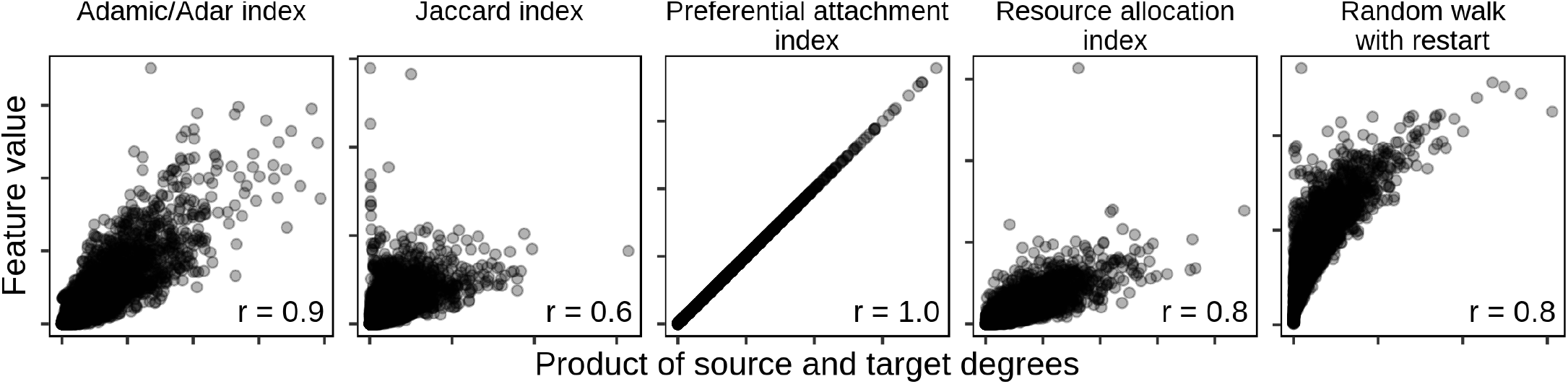
Common edge-prediction metrics correlate with node degree. Five common edge-prediction features (Supplemental table 2) are correlated with node degree on the STRING PPI network [18]. All five features show a positive relationship with degree, though the magnitude of this correlation is highly variable. The preferential attachment index is understandably perfectly correlated because it is equal to the product of source and target degree. Each panel indicates the Pearson correlation (“r”) between feature and degree in the lower right corner.

We used two permutation-derived null values to evaluate reconstruction and contextualize performance. First, the performance of the edge prior was compared to determine the performance attributable to the degree sequence of the PPI network. The first comparison gave insight into the ability of the PPI network to be reconstructed by degree. Second, the five edge prediction features were computed on 100 permuted networks and used to reconstruct the unpermuted network. Each permuted network corresponded to AUROC values quantifying the performances of features computed on it. The second comparison gave insight into the performance of each feature if the feature was only capturing degree.

The edge prior encapsulates nonspecific predictions due to degree, and it reconstructed the PPI network with an AUROC of 0.797 (dotted red line in Figure 8). In the second comparison, edge prediction features computed on permuted networks had performance equal or lower to their performances on the unpermuted networks. This indicated that four out of five edge prediction features discern more than node degree for the prediction task. The preferential attachment index is the product of source and target degree, and its performance did not differ from the edge prior or the feature’s performance when computed on permuted networks.

**Figure 8:**
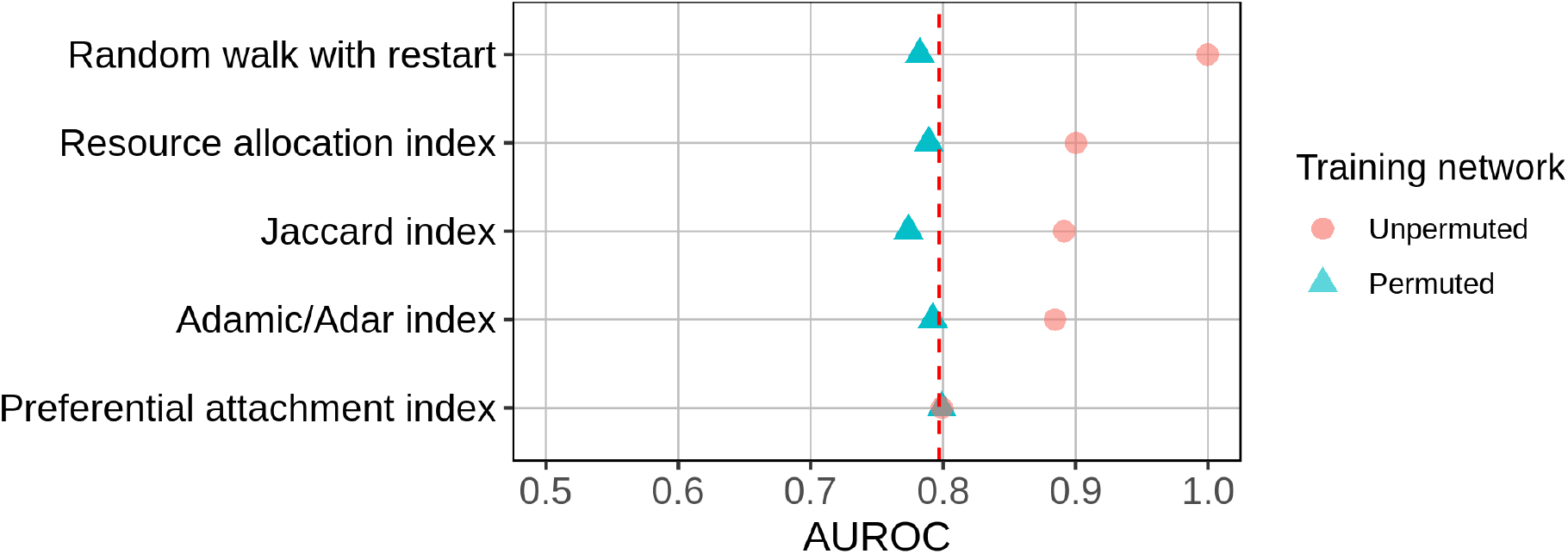
Identifying the fraction of a metric’s performance resulting from degree alone. Network reconstruction performances by five edge prediction features. Dotted red line indicates performance of the edge prior. Each feature was computed on both the unpermuted and 100 permutations of the STRING PPI network.

**Figure 9:**
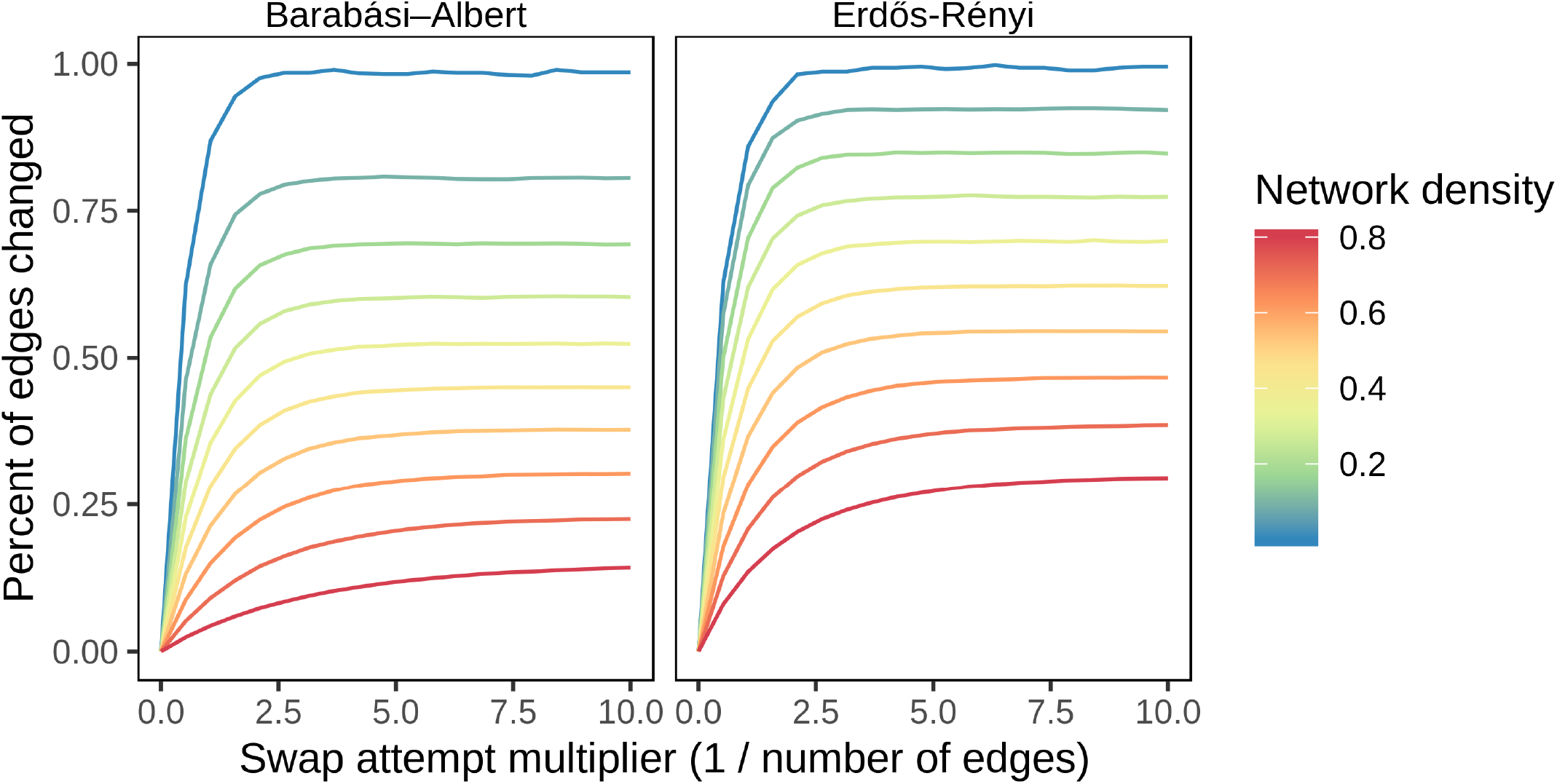
Higher density networks have lower asymptotic fractions of edges swapped and take more attempts to reach these values. The Barabási–Albert model produces scale-free random graphs, while Erdős–Rényi generates random graphs where all edges are equally likely.

This comparison quantified the performance of degree toward the prediction task and assessed degree’s effect on five edge prediction features. The edge prior provided the baseline level of performance attributable to degree alone. Comparing the performances on permuted networks to the performance of the edge prior reveals the extent to which a feature measures degree. Features whose performances on permuted networks were below that of the edge prior only imperfectly measured degree (eg: Jaccard index), whereas features whose performances equaled the edge prior completely captured degree (eg: preferential attachment index). Features can also capture information beyond degree, and our method can quantify this performance. For example, the superior performance on unpermuted networks relative to permuted networks indicated that RWR, resource allocation, Jaccard, and Adamic/Adar indices captured more than degree in this prediction task. These results aligned with the definitions of each feature and validated that our permutation framework accurately assessed reliance on degree.

## Discussion

We focus on edge prediction in biomedical networks. Our overall goal is to predict new edges with specificity, so that predictions reflect particular connectivity rather than generic node characteristics. Our permutation framework measures the predictive performance attributable to degree to provide a baseline expectation for edge pairs. We expect that non-specificity due to degree is not a unique property of biomedical networks. For example, if node A connects to nearly all other nodes in a network, predicting that all remaining nodes share an edge with node A will likely result in many correct—though nonspecific—predictions, regardless of the type of data contained in the network. Node degree should be accounted for to make correct predictions while being able to distinguish specific from nonspecific predictions. Prediction without reliance on node degree is challenging because many effective methods for edge prediction are correlated with degree (Figure 7).

The effects of node degree are obvious when edge prediction features are functions of degree. For example, the resource allocation index is the sum of inverse degree of common neighbors between source and target nodes (in the symmetric case), while preferential attachment is the product of source and target degree [26,27]. However, because many other edge prediction methods are not explicitly degree-based, it is important to have a general method for comparing the effects of node degree on edge prediction methods.

We developed a permutation framework to quantify the edge probability due to degree. We term this probability the “edge prior”, and we have identified two applications. First, a probability associated with every node pair can be treated as a classification score. Ordering these scores provides an assessment of performance based solely on degree, which can be used as a baseline for other classifiers. Second, node pair probabilities can be used to adjust edge prediction features depending on the task. If degree is a desired feature, then the edge prior can be treated like a Bayesian prior probability. Alternatively, if degree is not a desired feature, then the edge prior can be used to calibrate features and thus potentially enhance predictive specificity.

Figure 8 illustrates the utility of the edge prior and permutation framework for two purposes. First, it contextualizes feature performances relative to the baseline of nonspecific, degree-based predictions, quantified by the edge prior. Degree has varying utility for different edge prediction tasks. The edge prior’s performance on a task quantifies the utility of degree toward the task. This comparison is useful because specific predictions (based on more than degree alone) are more valuable for some applications than nonspecific ones and because degree can be an expression of bias in many real-world networks.

Second, Figure 8 compares five edge prediction features computed on and unpermuted networks. This comparison identified the fraction of each feature’s performance attributable to degree. Some features, such as the preferential attachment index, perfectly and exclusively measure degree. The Adamic/Adar index also almost completely captures degree because its performances from permuted networks are nearly at the performance of the edge prior. However, the Adamic/Adar index had much higher performance when computed on the unpermuted network, indicating that it also extracts higher-order information. This analysis, enabled by network permutation, measured the extent to which features rely on degree for a specific prediction task by assessing performance beyond the degree-based, nonspecific baseline.

## Conclusion

We developed a network permutation framework and open source software implementation that quantifies the probability of edge existence due to degree and can assess the fraction of feature performance attributable to degree. We demonstrated the superiority of the edge prior over other degree-based features for quantifying the effect of degree on the probability of edge existence. The XSwap methods and software provide a context for evaluating edge prediction methods and specific predictions for reliance on degree and, therefore, nonspecificity. Network edge prediction is a common task in biological and biomedical research, and it can be greatly influenced by degree. Degree should be considered directly in prediction approaches to avoid making nonspecific or trivial predictions due to degree imbalance or bias. A careful accounting of degree’s effects enables contextualized model evaluation and can help to quantify nonspecificity in biomedical network edge prediction.

## Acknowledgments

The authors thank Blair Sullivan for her feedback on a draft of the manuscript.

## Supplemental information

### XSwap parameter settings for network types

**Table 1:**
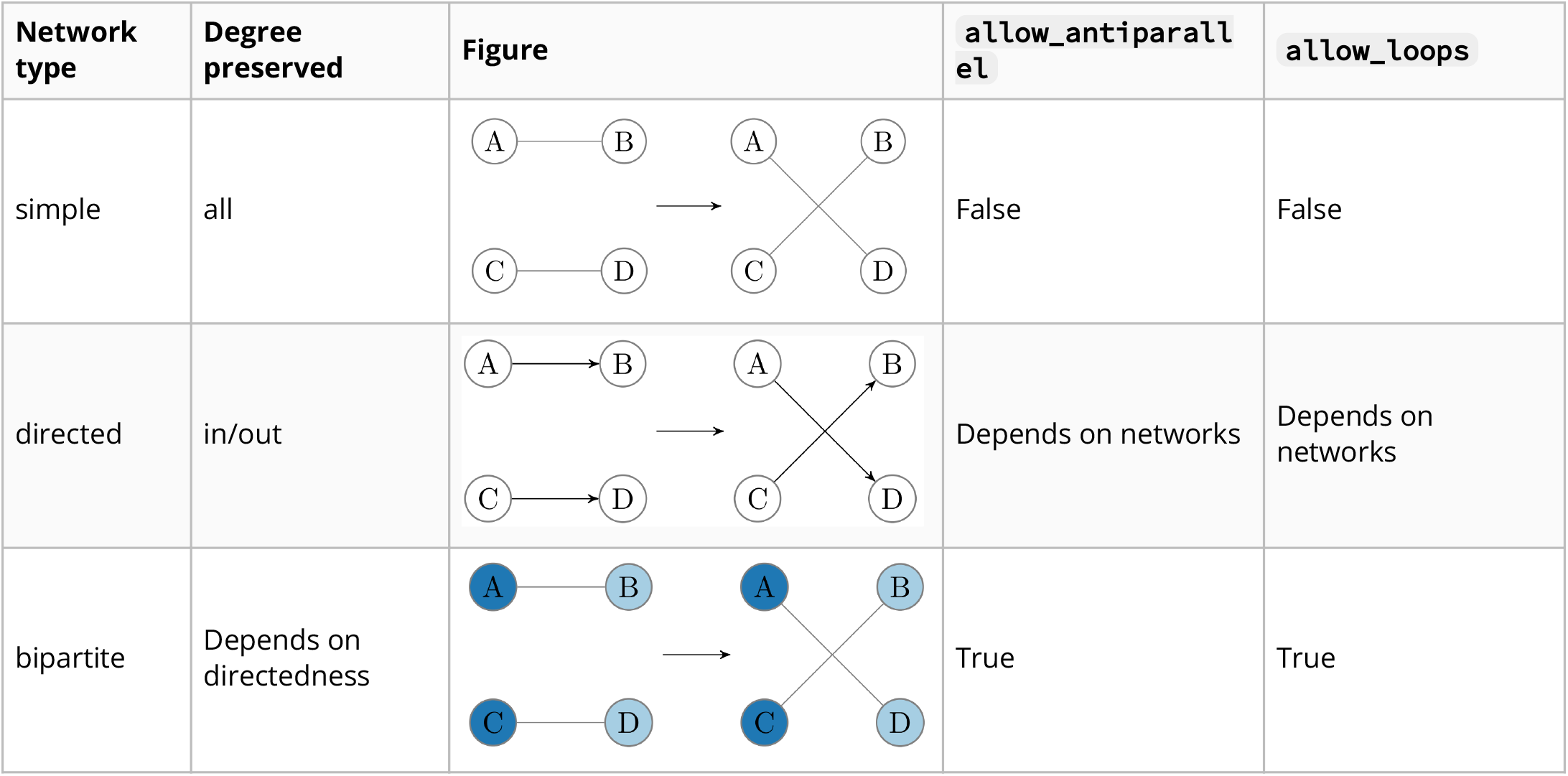
Applications of the modified XSwap algorithm to various network types with appropriate parameter choices. For simple networks, each node’s degree is preserved. For bipartite networks, each node’s number of connections to the other part is preserved, and the partite sets (node class memberships) are preserved. For directed networks, each nodes’ in- and out-degrees are preserved, though parameter choices depend on the network being permuted. Some directed networks can include antiparallel edges or loops while others do not.

| Network type | Degree preserved | Figure | allow_antiparallel | allow_loops |
| --- | --- | --- | --- | --- |
| simple | all |  | False | False |
| directed | in/out |  | Depends on networks | Depends on networks |
| bipartite | Depends on directedness |  | True | True |

### Performance of the XSwap algorithm

The performance of the XSwap algorithm depends on a number of network properties. We define network density to be the number of edges divided by the number of potential edges. Increasing network density lowers the asymptotic fraction of edges changed, as greater density prevents the algorithm from removing certain edges. Random graphs generated with a preferential attachment mechanism (via Barabási–Albert) can have a lower fraction of their edges swapped, asymptotically, as compared to uniform random graphs (via Erdős–Rényi).

### Approximate edge prior

To approximate the edge prior, we began by making two simplifications. First, we assumed independence between node pairs. This assumption does not actually hold for the XSwap algorithm, though it is a reasonable simplification for large, sparse networks. Second, we assumed that the XSwap process is stationary. This assumption also does not actually hold, but it was made because it significantly simplifies the problem. A single node pair has two possible states, “edge” and “no edge”. These states are not transient, and they are not periodic so long as more than one possible swap exists in the network. In almost all cases, then, our simplified model of the algorithm gives the state of a node pair as an ergodic process, independent of other node pairs.

Let *A*_*i*_,_*j*_ represent the existence of edge (*i, j*) For a given node pair, (*i, j*), then, let *q*_*i*_,_*j*_ represent the transition probability from the “no edge” state to the “edge” state in one successful iteration of the XSwap algorithm. Let *r*_*i*_,_*j*_ represent the probability of the opposite transition (“edge” to “no edge”) in one successful iteration. With “no edge” represented as [**1, 0**]^*T*^ and “edge” represented as [**0, 1**]^*T*^, the transition matrix, *P*, is given by the following:

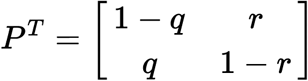

The stationary distribution of this system should correspond to the distribution when the number of swaps goes to infinity. It can be found by computing the eigenvectors of the system, as we know that the stationary distribution vector, **v** satisfies ***P*** ^***T***^ **v = v**. The eigenvector **v**, normalized to sum to 1 as a probability vector, is given by

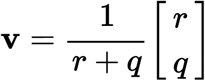

The asymptotic edge probability is therefore

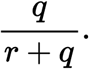

Since node pairs are being treated as independent, the probability of an edge being created in one successful iteration, given that the edge does not currently exist, is the ratio of the number of edge choices involving nodes *i* and *j* to the total number of possible swaps, *S*. Let *d*(*u*_*i*_) represent the degree of source node *i* and *d*(*v*_*j*_) represent the degree of target node *j*.

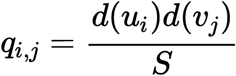

Similarly, the probability of an edge being eliminated in one iteration is the ratio of the number of edge choices involving (*i, j*) and any other valid edge to the total number of possible swaps. Let *m* be the total number of edges in the network.

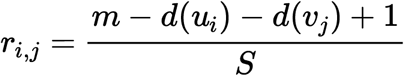

The approximate edge prior is, therefore,

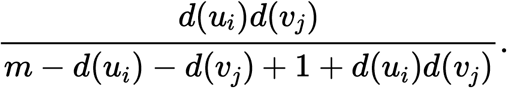

Unfortunately, we found that the above edge prior approximation is a poor approximation in many cases. We found that the following modified form (introduced in Methods) affords a superior approximation:

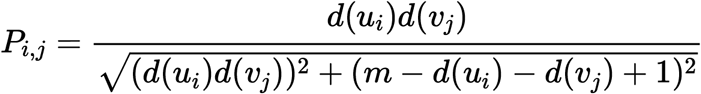

Interestingly, this expression can be derived by normalizing the eigenvector v to be a unit vector in the 2-norm instead of the 1-norm; that is, we use the value 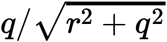 instead of *q*/(*r* + *q*). Because the modified form of the approximation offers a much superior fit to the data, we chose to include only the modified version in the released Python package, and we used the modified form throughout our analysis.

### Networks used for comparison

| Data | Network | Nodes | Edges |
| --- | --- | --- | --- |
| Hetionet | AdG | Source: 402, Target: 20945 | 102240 |
|  | AeG | Source: 402, Target: 20945 | 526407 |
|  | AID | Source: 402, Target: 137 | 3602 |
|  | AuG | Source: 402, Target: 20945 | 97848 |
|  | BPpG | Source: 11381, Target: 20945 | 559504 |
|  | CCpG | Source: 1391, Target: 20945 | 73566 |
|  | CbG | Source: 1552, Target: 20945 | 11571 |
|  | CcSE | Source: 1552, Target: 5734 | 138944 |
|  | CdG | Source: 1552, Target: 20945 | 21102 |
|  | CrC | 1552 | 6486 |
|  | CuG | Source: 1552, Target: 20945 | 18756 |
|  | DaG | Source: 137, Target: 20945 | 12623 |
|  | DdG | Source: 137, Target: 20945 | 7623 |
|  | DpS | Source: 137, Target: 438 | 3357 |
|  | DuG | Source: 137, Target: 20945 | 7731 |
|  | GuG | 20945 | 265672 |
|  | GcG | 20945 | 61690 |
|  | GiG | 20945 | 147164 |
|  | GpMF | Source: 20945, Target: 2884 | 97222 |
|  | GpPW | Source: 20945, Target: 1822 | 84372 |
| PPI | Sampled | 3992 | 255522 |
|  | Literature | 3992 | 364743 |
|  | Systematic | 3916 | 12913 |
| bioRxiv | Sampled | 4587 | 30686 |
|  | <2018 | 4615 | 43691 |
|  | All time | 4615 | 44963 |
| TF-TG | Sampled | Source: 142, Target: 1396 | 2689 |
|  | Literature | Source: 144, Target: 1406 | 3496 |
|  | Systematic | Source: 144, Target: 1417 | 29177 |

### Edge prediction features

In the table that follows, let *k*(*u*) denote the set of neighbors of node *u*. Let **A** represent the normalized Laplacian adjacency matrix, and let *y*_*u*_ be a vector with all ones except for a one in the *u*-th position. *x* For a directed graph, let ***A***(***u***) denote the set of nodes that node *u* points to and ***D***(***u***) the set of nodes that point to *u*. All definitions that follow are the score between nodes ***u*** and ***v***.

**Table 2:**
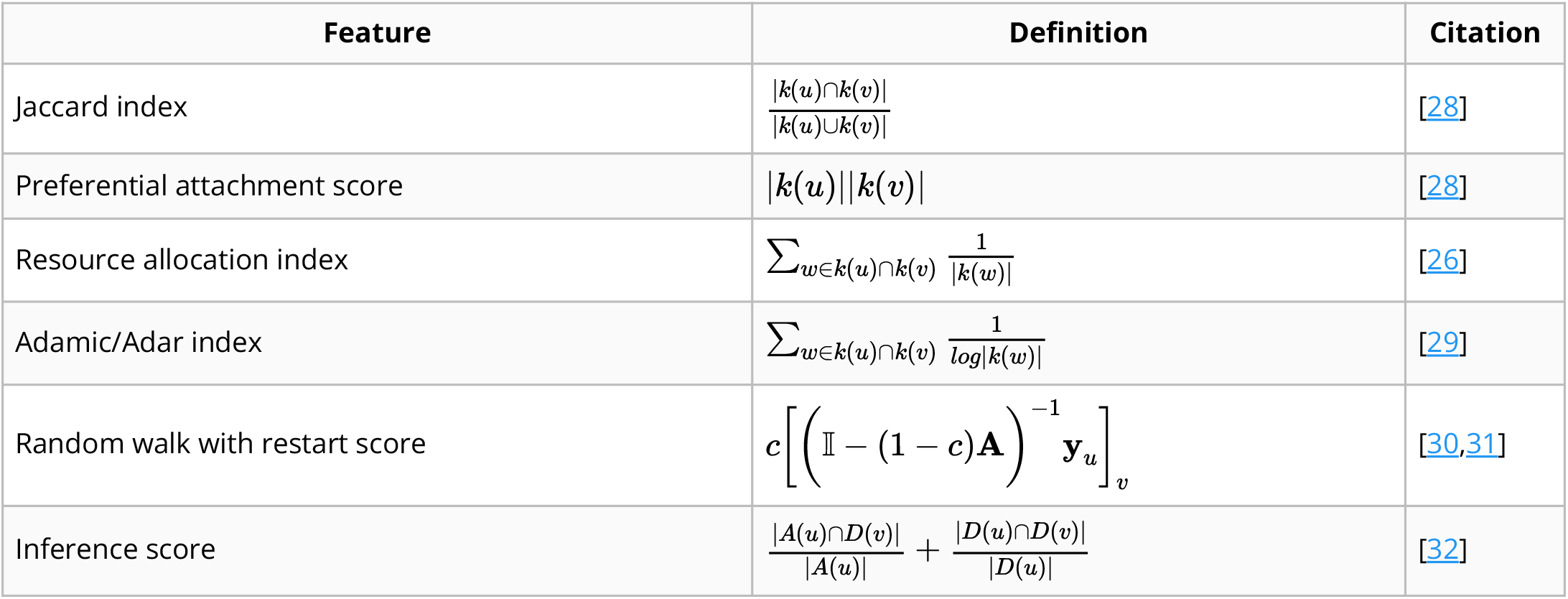
Edge prediction features.

